# Manuscript 101: a data-driven writing exercise for beginning scientists

**DOI:** 10.1101/139204

**Authors:** Michael A. Halbisen, Amy Ralston

## Abstract

Learning to write a scientific manuscript is one of the most important and rewarding scientific training experiences, yet most young scientists only embark on this experience relatively late in graduate school, after gathering sufficient data in the lab. Yet, familiarity with the process of writing a scientific manuscript and receiving peer reviews, often leads to a more focused and driven experimental approach. To jump-start this training, we developed a protocol for teaching manuscript writing and reviewing in the classroom, appropriate for new graduate or upper-level undergraduate students of developmental biology. First, students are provided one of four cartoon data sets, which are focused on genetic models of animal development. Students are instructed to use their creativity to convert evidence into argument, and then to integrate their interpretations into a manuscript, including an illustrated, mechanistic model figure. After student manuscripts are submitted, manuscripts are redacted and distributed to classmates for peer review. Here, we present our cartoon datasets, homework instructions, and grading rubrics as a new resource for the scientific community. We also describe methods for developing new datasets so that instructors can adapt this activity to other disciplines. Our data-driven manuscript writing exercise, as well as the formative and summative assessments resulting from the peer review, enables students to learn fundamental concepts in developmental genetics. In addition, students practice essential skills of scientific communication, including arguing from evidence, developing and testing models, the unique conventions of scientific writing, and the joys of scientific story telling.

## Introduction

Manuscripts are one of the main products of academic scientific research. Accordingly, learning how to effectively package observations and interpretations into a compelling, logical, and mechanistic story is a major goal of graduate training. However, for most graduate students, lessons in manuscript writing become available relatively late in their training – after students have generated sufficient data. Arguably, an intimate knowledge of the manuscript-writing process should precede and drive the experimental approach. We propose that modeling the processes of writing and reviewing a manuscript in the classroom can accelerate the development of skills that are essential for independent laboratory study (Table 1).

**Table 1.** Manuscript 101 Concepts and Skills. This two-part homework assignment models manuscript writing and reviewing. Our specific data sets were designed for a first-year graduate course in developmental genetics, but could be adapted to other disciplines in biology, and to include more or less advanced students. The goals of the homework are to expose students to the following conceptual biological topics and transferrable scientific skills (Hunter and Metevier, 2011):

Biological Concepts
- Gaining an understanding of genetic approaches used in model organisms
- Using multiple methods for evaluating gene expression analysis
- Using positive and negative controls
- Identifying multiple biological interpretations that are consistent with a given experimental observation

Transferrable Scientific Skills
- Arguing from evidence
- Structuring a scientific manuscript
- Employing the creative process of scientific storytelling
- Integrating multiple lines of evidence to conceive a mechanistic model
- Stating and testing predictions of an integrated model
- Understanding/anticipating criticism as an opportunity to strengthen the evidence

To address the need for early exposure to manuscript writing, we developed a two-part homework assignment that enables beginning graduate or upper-level undergraduate students to experience the process of writing and submitting a manuscript (Fig. 1). This assignment, which we call Manuscript 101, was tested and improved, after written student assessment, over the course of four offerings of a first year graduate level course in developmental genetics (n = 43 students). For Homework 1, each student is provided with the manuscript-writing instructions (Fig. S1) as well as one of four sets of cartoon figures (Fig. 2 and Fig. S2-5). The figures provided are cartoon data, and are provided in unsequenced order. Students are instructed to organize their figures in an order that tells the best story, to interpret the data, to draw a model figure illustrating their proposed molecular mechanism, and to write a manuscript using all of the figures. For the manuscript, students write Title, Abstract, Introduction, Results, and Discussion sections, according to the *Author Guidelines* provided in the Instructions for Homework 1 (Fig. S1). In the Discussion section, students are encouraged to provide alternative interpretations of their data, and to propose experiments that would discriminate among the possible interpretations. Students are also encouraged to state the predictions of their model, and to propose future experiments that would test these predictions. Thus, Manuscript 101 exposes students to the structure and organization of a typical scientific manuscript, but also provides students with a first-hand opportunity to experience the creative aspects of scientific story-telling and exploration.

**Figure 1.**
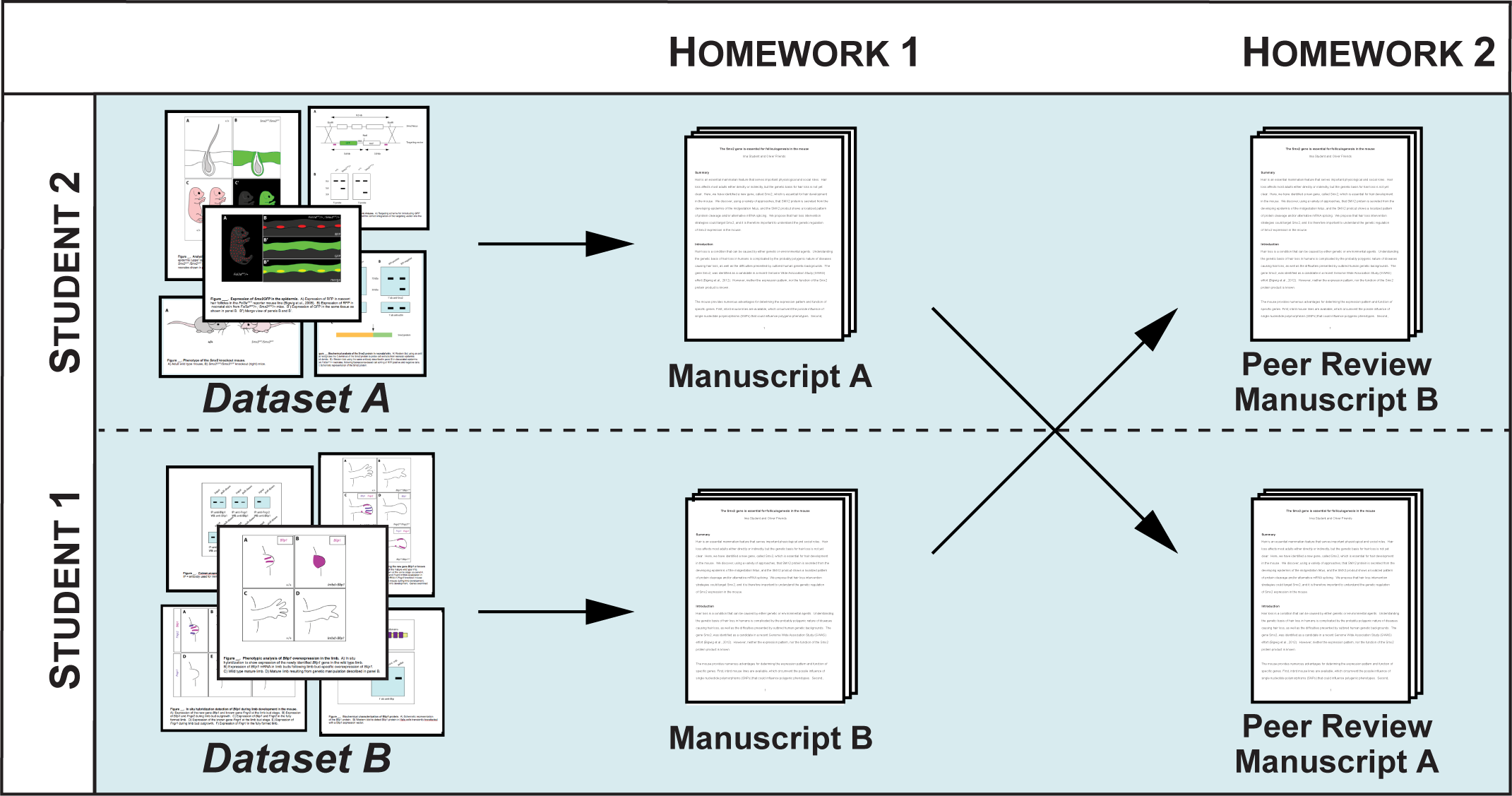
Overview of the two-part Manuscript 101 homework. For homework 1, students are each assigned a cartoon dataset (Fig. S2-S5), and the instructor explains the instructions, which are provided as lecture slides (Fig. S1). Students write their manuscripts and submit these to the instructor, who serves as journal editor and an additional reviewer during the mock peer review of homework 2. Instructions for Homework 2 are included as lecture slides (Fig. S6).

**Figure 2.**
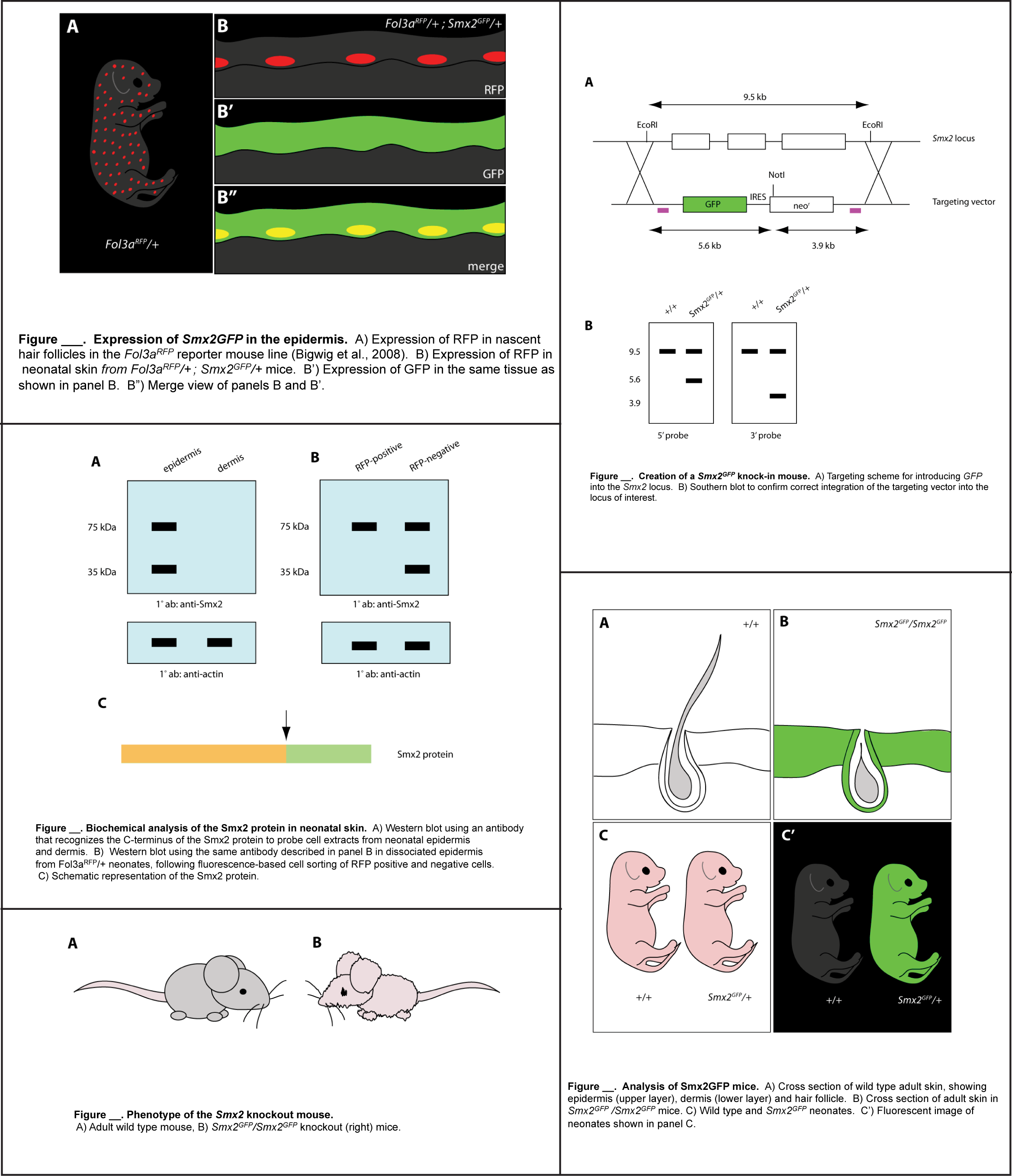
Example of Datasets used for manuscript writing homework. Students are provided one of the four datasets (Fig. S2-S5), shown here is Dataset B (Fig. S2). Figure panels are not in order, but are to be ordered and labeled by students according to the flow of their manuscript. Basic legends are provided, and students are encouraged to edit as needed. Students interpret the figures in the Results section of their manuscripts, and include a final model figure, which draws their interpretations into a mechanistic and testable model.

The manuscripts are submitted to the instructor, who acts as journal editor in Homework 2, the mock manuscript review. During Homework 2, each student is asked to provide an anonymous review of a manuscript generated by a peer in Homework 1, according to the *Reviewer Guidelines* provided in the Instructions for Homework 2 (Fig. S6). The instructor may also choose to provide a review of each submitted manuscript.

Therefore, Homework 2 is both an assessment of the manuscript writing assignment, as well as an experiential opportunity in critical evaluation of scientific manuscripts. At the end of Homework 2, each student will receive the review(s) of his or her manuscript, as well as a formal assessment of both homework assignments from the instructor. Below, we provide the teaching tools (Table 2), and framework for implementing this practical exercise.

**Table 2.** Teaching Tools. The following resources are included for this manuscript writing and reviewing modeling activity. After in-class introduction, students pursue the activity outside of lecture.

1. Homework 1 Instructions (Fig. S1)
2. Sample Data Sets (Fig. S2-S5)
3. Homework 2 Instructions (Fig. S6)
4. Evaluation Tools (see Table 3)

## Results and Discussion

### 1. Homework How-To

Instructions for Homework 1 and 2 (Fig. S1 and S6) are provided to each student, along with a cartoon data set (Fig. S2-S5). The goal of Homework 1 is to produce a manuscript, which will be peer-reviewed as Homework 2. It is important that students receive instructions for Homework 2 at the same time that they receive the instructions for Homework 1 because this will enable students to understand the assessment criteria for Homework 1. Homework 1 instructions include an overview of the writing exercise, as well as section-by-section manuscript guidelines to clarify content, format, and word limits. Depending on the class size, multiple students may receive the same data set. In this case, students are instructed that they are permitted to work together to interpret their data, as long as intellectual contributions are acknowledged by coauthorship. However, the instructions also specify that each student is to submit an original, independently written manuscript.

When all students have submitted Homework 1, each manuscript is assigned an anonymous identifier that does not include student names. Each manuscript is then redacted to protect student identity, and then each manuscript is assigned to another student as Homework 2. Ideally, the data sets used by author and peer reviewer should differ. To help keep track of student names, their data sets, and anonymous identifiers, we provide an organizational spreadsheet (Table S1).

The goal of Homework 2 is to provide students with a realistic peer-review experience, from the perspective of the author and the reviewer. Homework 2 instructions include clear criteria for evaluating a manuscript. Students should also understand that Homework 2 cannot be assigned until each student has submitted Homework 1. Prior to each homework assignment, it is beneficial to review homework instructions (Fig. S1 and S6) during lecture, and to assess student familiarity of the structure and style of scientific writing (Table 3). Although most students will have received training in the reading and critical evaluation of scientific manuscripts, most have not yet engaged in the writing of manuscripts or manuscript reviews.

**Table 3.** Evaluation Tools. A. **Summative Student Assessment**

- Instructor’s organizational spreadsheet (Table S1)
- Homework 1 Scoring matrix (Table S2)
- Homework 2 Scoring Matrix (Table S3)
B. **Formative Activity Assessment**
  Questions to ask students before and after Homework 1
  - What is the purpose of each section of a scientific manuscript?
  - What topics are appropriate for each section of a scientific manuscript?
  - Is there only one way to tell a scientific story?
  - Is there only one way to interpret a given observation?
  - Once a mechanistic model is produced, how can it be tested?
  Questions to ask students before and after Homework 2
  - What is a typical organization of a review of an unpublished manuscript?
  - How should the review process change your approach to scientific writing?
  - What are some strategies for ensuring that a critical evaluation of a manuscript is constructive?

### 2. Homework Assessment

Tools for assessing student performance, as well as homework efficacy, are listed in Table 3. Logistically, it is helpful for the professor to create a spreadsheet to keep track of manuscript identifiers, reviewer assignments, and final grades. A sample organizational spreadsheet is provided (Table S1). For Homework 1, students receive two independent reviews of their manuscript – one written by an anonymous student peer, and one written by the instructor. If time permits, the instructor can also provide a more detailed assessment of Homework 1 by annotating each student’s manuscript.

In addition, students receive a grade for each of the two homework assignments. To facilitate grading, we have provided two scoring matrices (Tables S2 and S3), which closely track the expectations stated in the homework instructions. Results for the scoring matrices may be returned to the students, or summarized as comments. Typical assessment commentary includes a paragraph highlighting the major points that drove the grades for each assignment.

### 3. Designing New Data Sets

We have included sample data sets that were designed for a graduate level course in vertebrate developmental genetics. Manuscript 101 can be adapted to include additional model organisms and biological processes by devising new cartoon data sets. Here, we describe our approach to creating the cartoon data sets in order to provide a path for creating new data sets.

To devise our sample data sets, we used four criteria, which are generalizable to multiple biological disciplines. Our first criterion was that the cartoon data set should focus on processes and methods that have been described in lectures. In this way, the manuscript writing and reviewing homework can synergize with the didactic lesson. The second criterion was that each data set should highlight techniques for creating gain and loss of function (in genes or tissues) in order to test specific hypotheses. For example, our data sets focused on various model organism-specific approaches for altering gene expression. In this way, the homework teaches experimental approaches as well as concepts. The third criterion was that each data set should highlight standard approaches for evaluating experimental outcomes. For example, the endpoints of altered gene expression levels in our model data sets were embryo morphology and gene expression. Accordingly, our data sets included embryo morphology and gene expression data that would have been generated using microscopy, quantitative PCR, RNA-sequencing, *in situ* hybridization, immunofluorescence, flow cytometry, and western blotting approaches. Thus, the manuscript writing and reviewing homework is an opportunity to expose students to multiple experimental approaches for evaluating experimental endpoints. The final criterion used in creating our data sets was that several of our cartoon results could be interpreted in multiple ways. For example, a band shift on a gel could be due to alternative mRNA splicing or to protein cleavage. In this way, students are challenged to consider multiple alternative interpretations of an experimental test and to propose how they could discriminate between the possibilities in future experiments. New figures can be hand-drawn, or prepared using any computer illustration software.

## Conclusions

Manuscript 101 is designed to model some of the most important scholarly activities of working scientists, so that students can experience these activities early in their training. An additional benefit of this activity is that it exposes students to the creativity that is intrinsic to experimental science, as well as the elegant structure imposed by the scientific method and manuscript conventions. From the instructor’s perspective, it is fascinating to observe diverse models that arise from a singular set of figures, and rewarding to coach beginning scientists through some of their first forays in science communication.

## Supporting Figures

Figure S1. Instructions for Homework 1

Figure S2. Dataset A

Figure S3. Dataset B

Figure S4. Dataset C

Figure S5. Dataset D

Figure S6. Instructions for Homework 2

Table S1. Instructor’s organizational spreadsheet

Table S2. Homework 1 Scoring Matrix

Table S3. Homework 2 Scoring Matrix

## Homework 1 Instructions – Write your own manuscript!

1. **Overview**

a. **Goal**
b. **How-to**
c. **Assessment**
2. **Author Guidelines**

a. **Title**
b. **Abstract**
c. **Introduction**
d. **Results**
e. **Model figure with legend**
f. **Discussion**
g. **References**
h. **Format**
3. **Manuscript-writing resources**

### 1a. Goal: write a manuscript from cartoon data

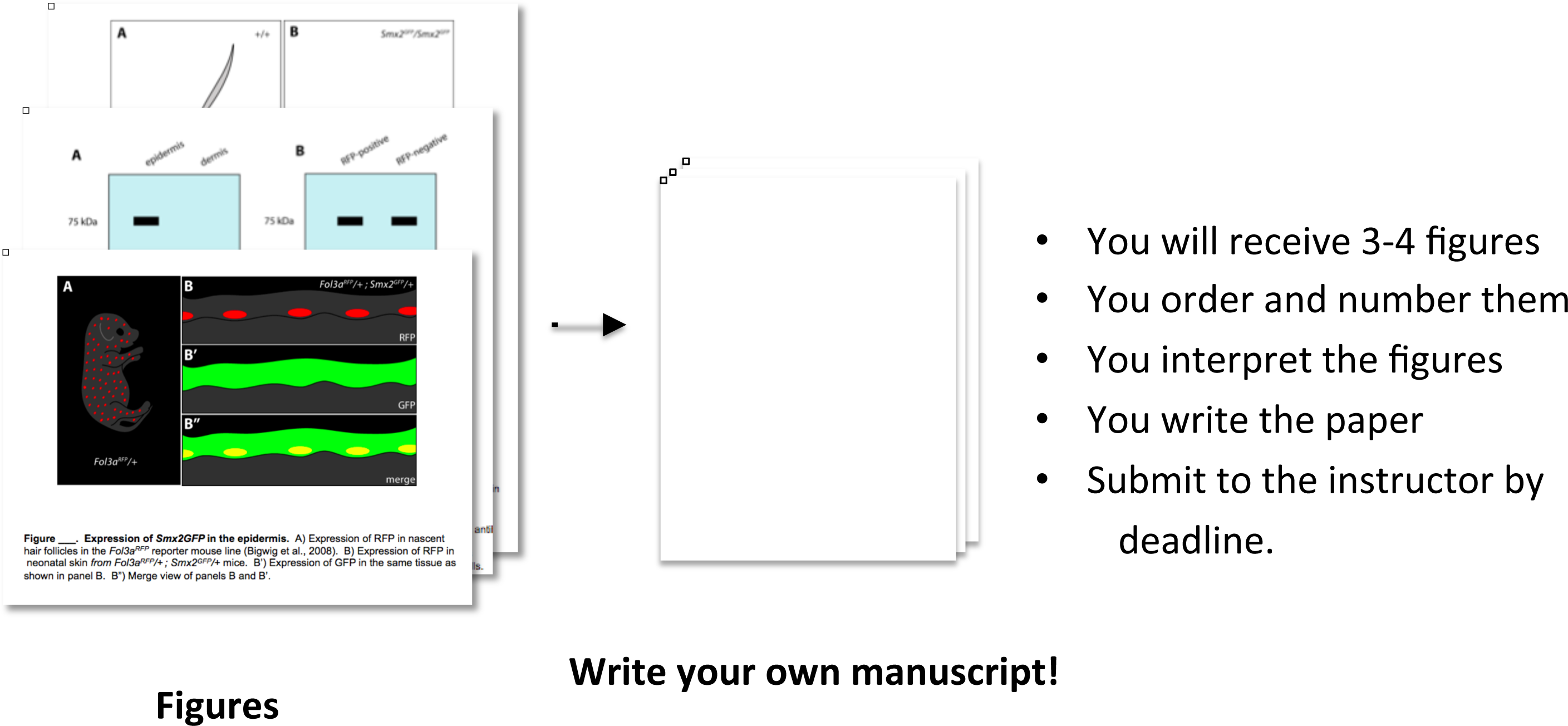

### 1b. How-To

- Follow *Author Guidelines (Section 2 of this talk)*
- You write: Title, Abstract, Introduction, Results, and Discussion sections
- You draw: a model figure and (write its legend)
- All writing must be your own
- Collaboration on interpreting figures is encouraged
- Collaborators must be listed as co-authors of your paper

Tips
1. Use your creativity to transform evidence into argument
2. Persuade your audience of your important new insight
3. Discover a new mechanism regula0ng the biological process at hand

### 1c. Assessments

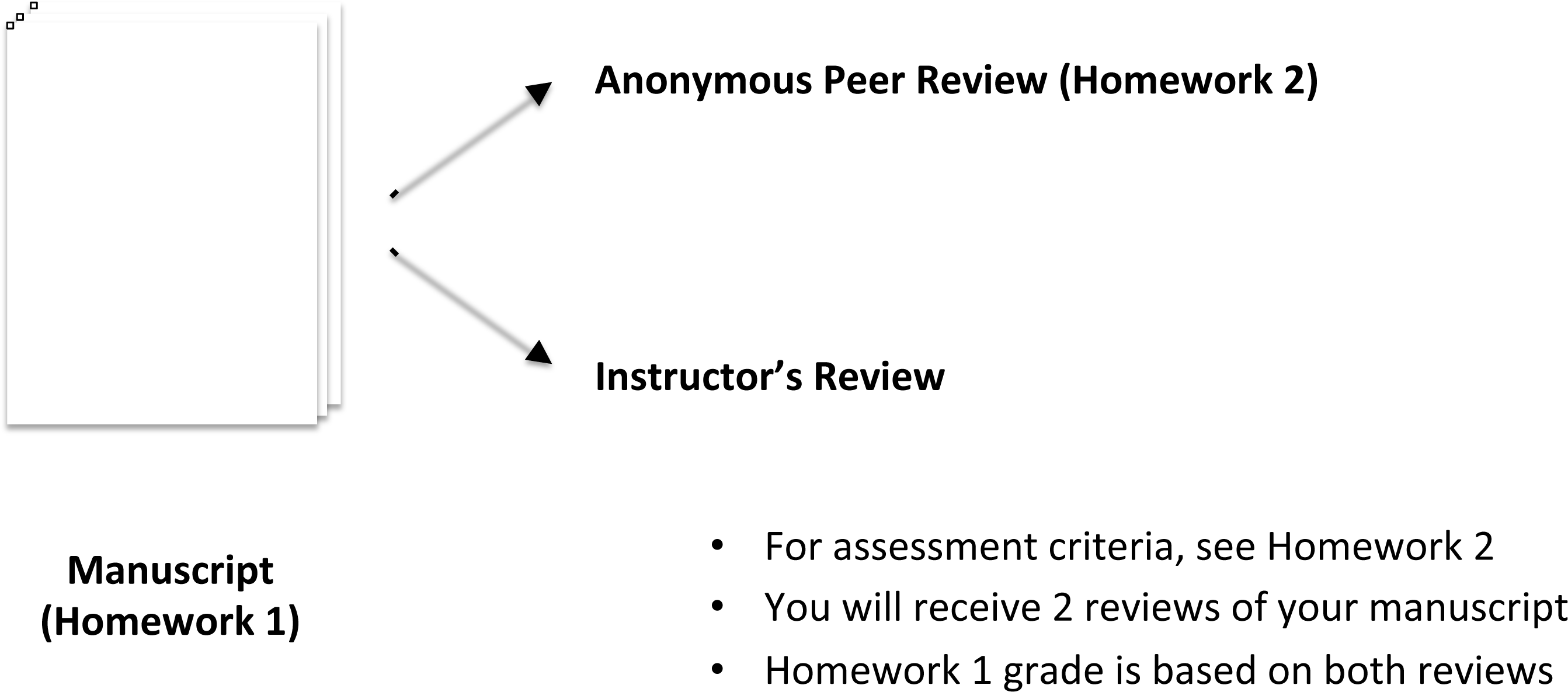

### 2a. Author Guidelines - Title

**Limit: 20 words**

- State the main lesson from the study

### 2b. Author Guidelines - Abstract

Limit: 200 words
- Pocket-sized synopsis of the paper
- Should entice the reader
- Should provide pithy summary of which new lessons are learned
- Should explain the **significance**:

1. Why would I have undertaken this study?
2. How could I get others excited about this study?
- Should explain the **discovery**:
  1. What do these pieces of data reveal about the process?
  2. What is the mechanism by which X molecule regulates Y process?
- Should explain the **novelty**:
  1. Why is this new?
  2. How does this differ from other studies?

### 2c. Author Guidelines - Introduction

Limit: 300 words
- Sets the stage for the story
- Provides background needed to understand the results
- Cite real papers for bonus points!
- Differs from Abstract and Discussion

- Abstract is the summary
- Discussion is the interpretation

### 2d. Author Guidelines - Results

Limit: 400 words per section
- Use section headings that summarize the key finding for each figure
- sections should follow the figures in sequence
- Refer to each figure panel parenthetically, the first time it arises
- Include in each section:

1. Brief rationale for each experiment (i.e., the hypothesis tested)
2. Brief description of the experimental approach, including controls
3. Your observations
4. Brief interpretation (i.e., do your observations support or reject the hypothesis?) *Remember that data can often be interpreted in multiple ways!*

### 2e. Author Guidelines - Model Figure and Legend

Legend Limit: 50 words
- A graphical summary of the mechanism you describe in the Discussion
- Legend should explain the model briefly and define colors/symbols used

### 2f. Author Guidelines - Discussion

Limit: 600 words
- Integrate all the interpretations into a mechanistic model
- The model is a new discovery about a biological process (e.g., the way that a protein regulates a developmental event)
- Refer to the model figure
- State the predictions of the model
- Propose experiments to test the predictions of the model
- Propose experiments to distinguish between alternate interpretations of results
- ***Avoid restating observations already explained in the Results section***

### 2g. References

No limit
- Cite real or fictinal papers throughout the text
- Use Endnote or other reference manager
- Use *Cell* reference style or similar

### 2h. Author Guidelines - Format

The Text
- 11 point Times New Roman or Arial
- Double-spaced
- 1-inch margins on all sides
- Page number, lower right hand corner
- Print to PDF

The Figures
- Number the figures and arrange in the order described in the text
- Edit legends if needed
- Print to PDF

Manuscript for Submission
- Merge Text and Figure PDFs, in that order
- Submit PDF of the complete manuscript to the instructor by email.

## 3. Other Resources

*Development* Author Instructions:

http://dev.biologists.org/site/misc/submissions.xhtml

Understanding the publishing process (Elsevier):

http://www.elsevier.com/journal-authors/publishing-process

**Figure ___.**
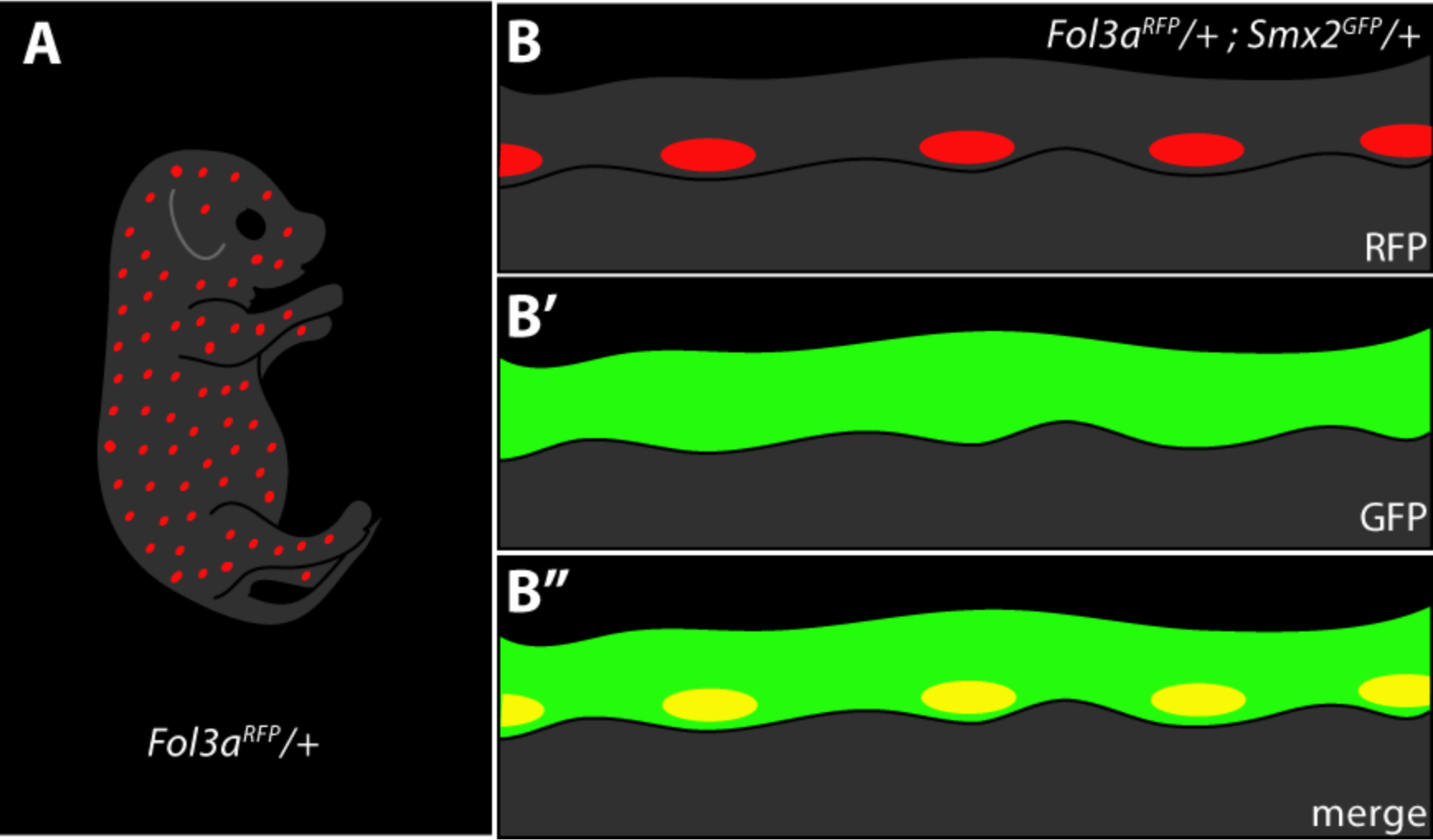
Expression of *Smx2GFP* in the epidermis. A) Expression of RFP in nascent hair follicles in the *Fol3a*^*RFP*^ reporter mouse line (Bigwig et al., 2008). B) Expression of RFP in neonatal skin from *Fol3a^RFP^*/+; *Smx2*^*GFP*^/+ mice. B’) Expression of GFP in the same tissue as shown in panel B. B′′) Merge view of panels B and B′.

**Figure __.**
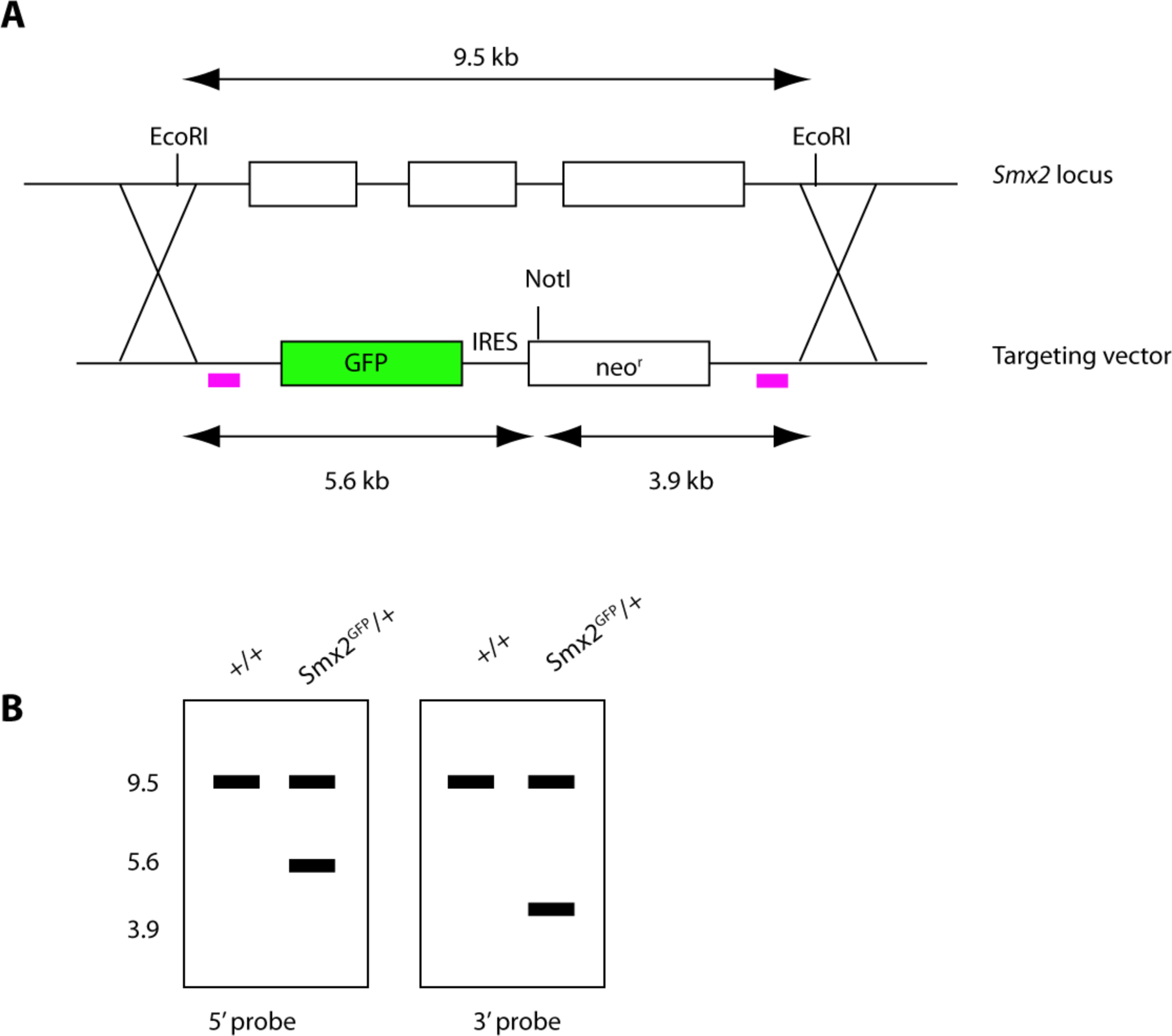
Creation of a *Smx2*^*GFP*^ knock-in mouse. A) Targeting scheme for introducing *GFP* into the *Smx2* locus. B) Southern blot to confirm correct integration of the targeting vector into the locus of interest.

**Figure __.**
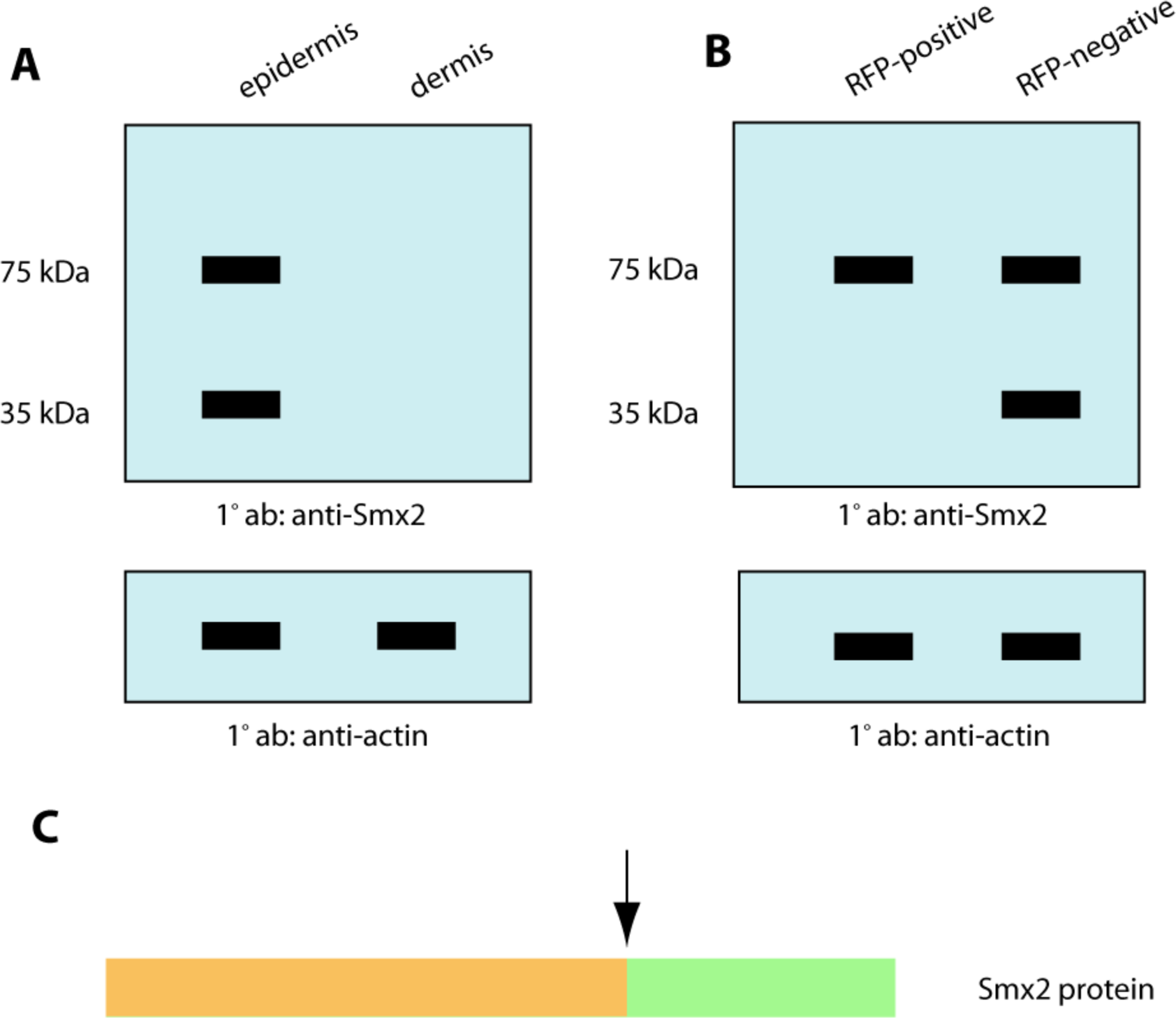
Biochemical analysis of the Smx2 protein in neonatal skin. A) Western blot using an antibody that recognizes the C-terminus of the Smx2 protein to probe cell extracts from neonatal epidermis and dermis. B) Western blot using the same antibody described in panel B in dissociated epidermis from Fol3a^RFP^/+ neonates, following fluorescence-based cell sorting of RFP positive and negative cells. C) Schematic representation of the Smx2 protein.

**Figure __.**
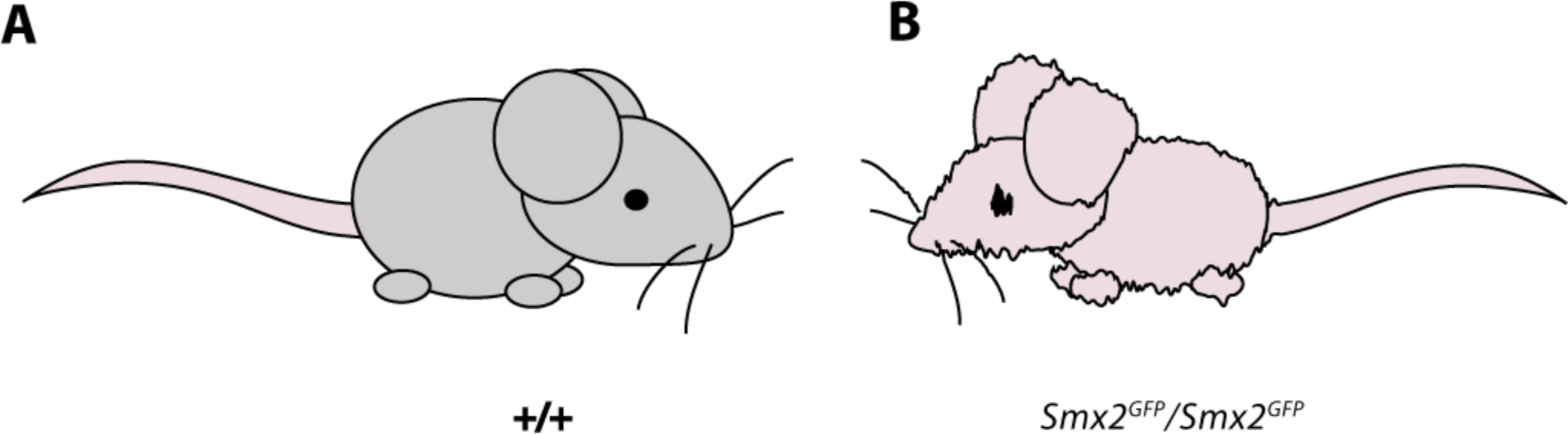
Phenotype of the *Smx2* knockout mouse. A) Adult wild type mouse, B) *Smx2*^*GFP*^/*Smx2^GFP^* knockout (right) mice.

**Figure __.**
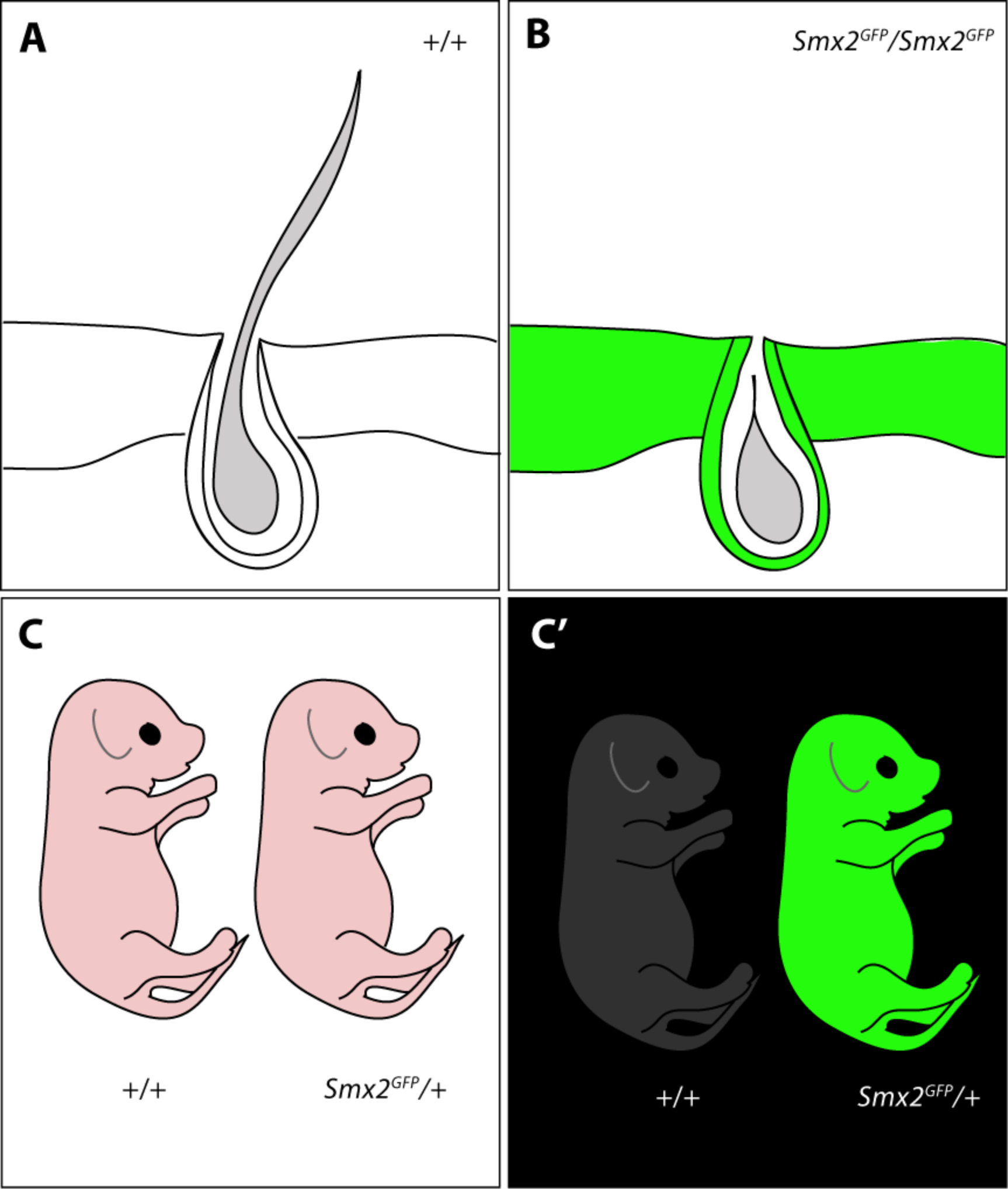
Analysis of Smx2GFP mice. A) Cross section of wild type adult skin, showing epidermis (upper layer), dermis (lower layer) and hair follicle. B) Cross section of adult skin in *Smx2*^*GFP*^/*Smx2*^*GFP*^ mice. C) Wild type and *Smx2*^*GFP*^ neonates. C’) Fluorescent image of neonates shown in panel C.

**Figure __.**
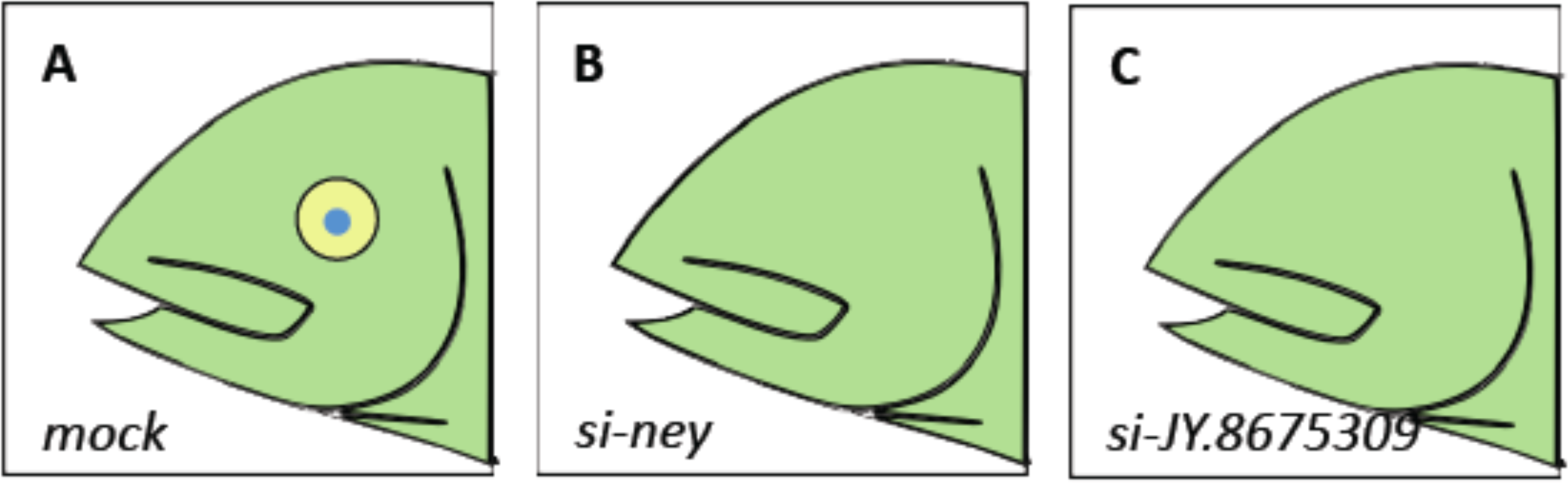
Phenotypes resulting from siRNA treatment of larval proto-eyes. A) Typical phenotype resulting from embryo injection with scrambled siRNAs. B) Phenotype resulting from injection of *ney* siRNA. C) Phenotype resulting from injection of siRNA specific to novel gene JY.8675309.

**Figure __.**
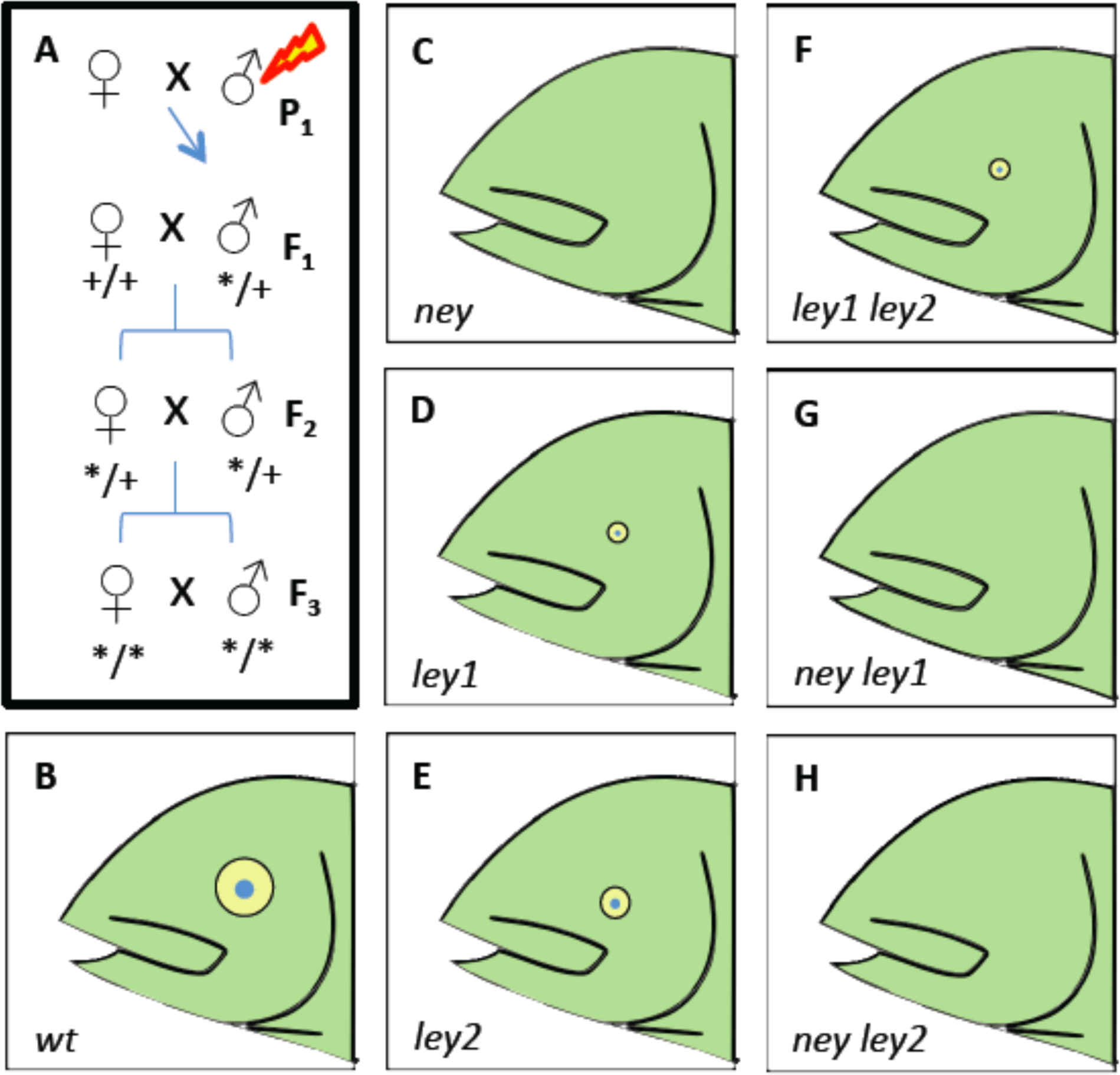
Phenotypes of mutants recovered from F3 mutagenic screen. (A) Mutagenesis and genetic screen strategy. (B) wild type (C) *no eye mutant* (D) *little eye 1 mutant (E) little eye 2 mutant (F) ley1 ley2 double mutant* (G) *ney ley1 double mutant (H) ney ley2 double mutant.*

**Figure __.**
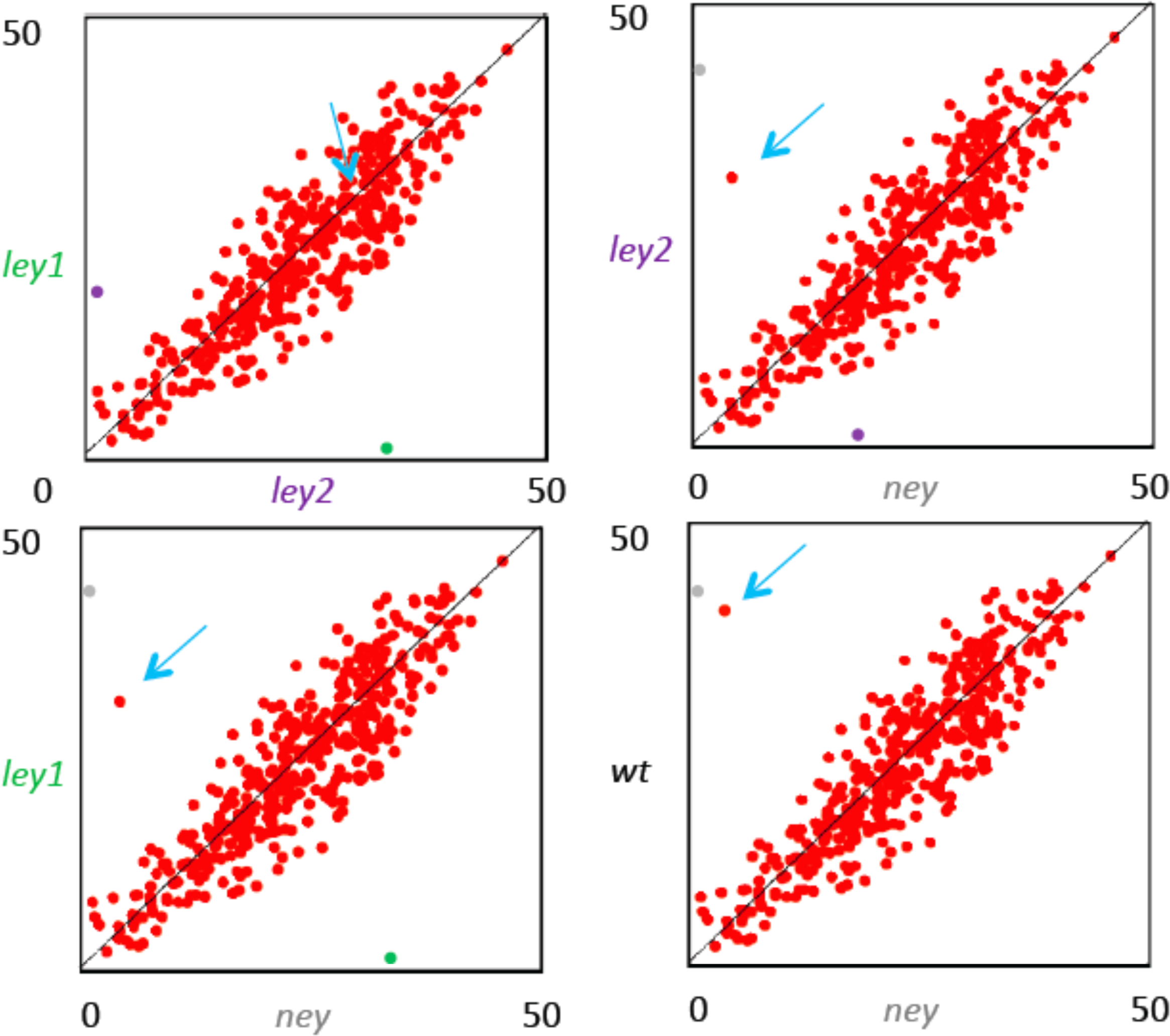
Correlations between mutant and wild type larval transcript levels. In each panel a dot is color-coded probes to match the gene on *x or y axis. The blue arrow* indicates an unknown transcript (JY.8675309).

**Figure __.**
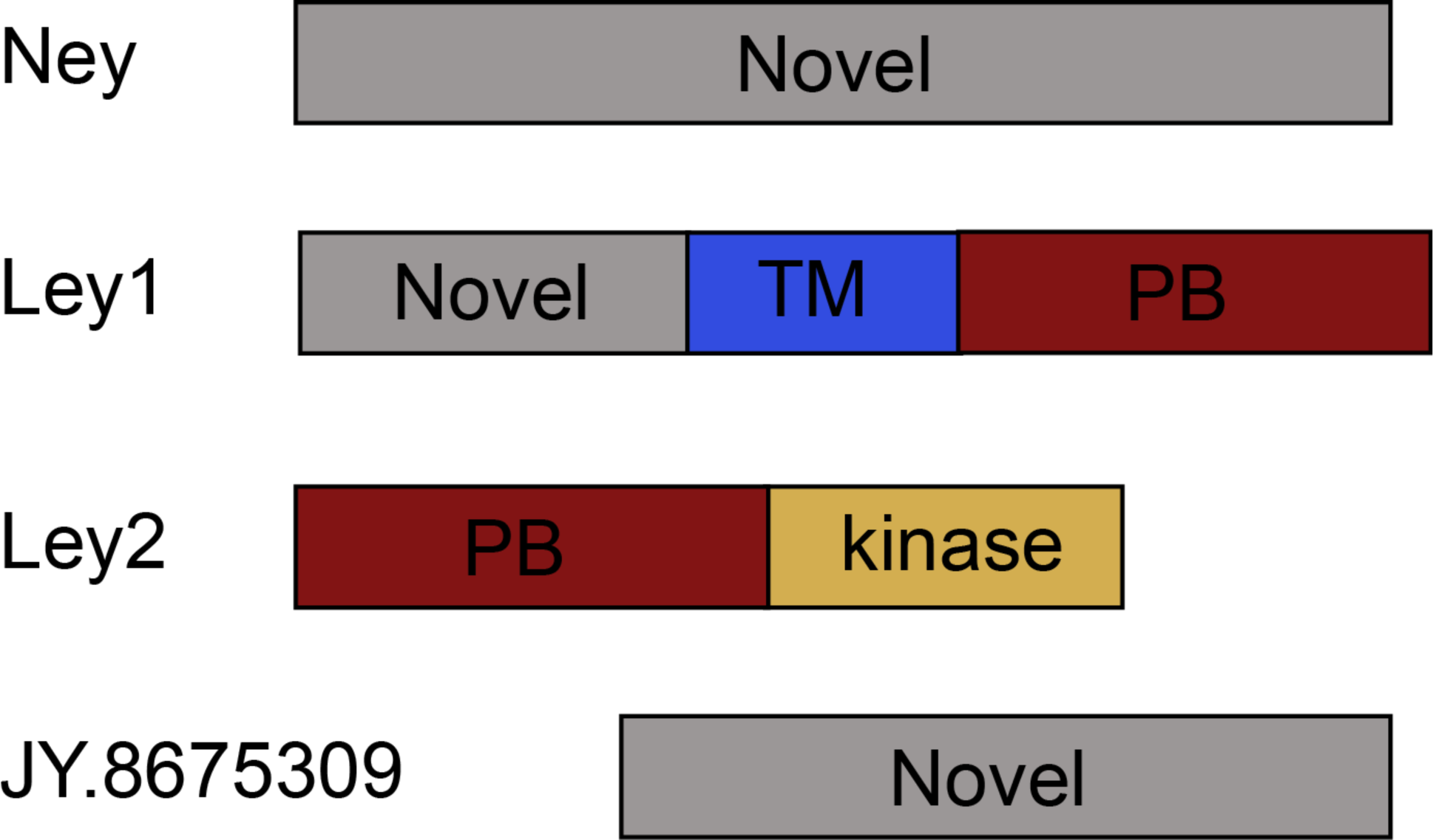
Predicted domains of proteins encoded by newly identified genes. Novel = no known funciton, TM = putative transmembrane domain, PB = putative protein binding domain, kinase = putative kinase activity.

**Figure __.**
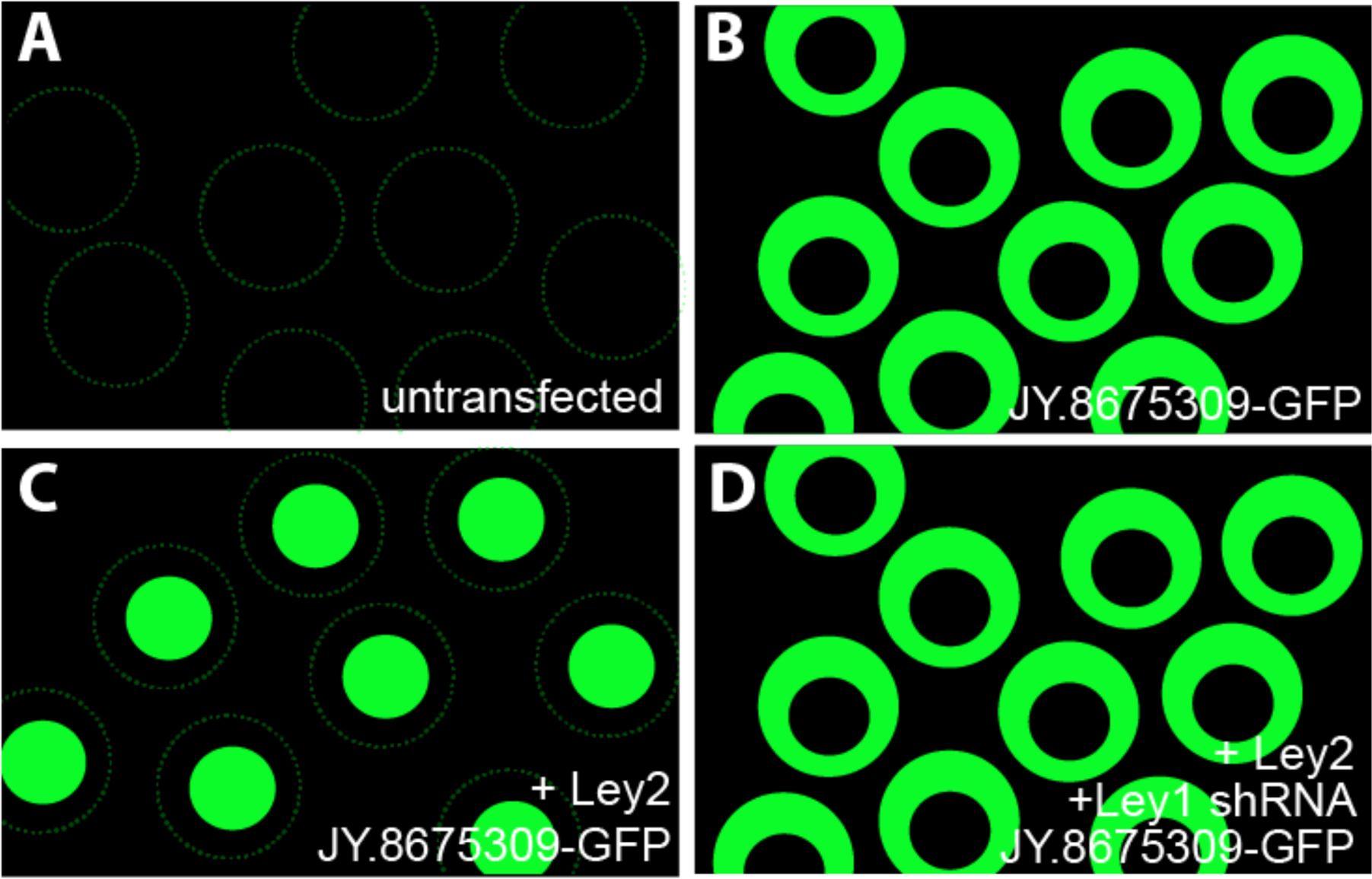
Subcellular localization of JY.867 in cultured cell line. A) Background fluorescence in untransfected cells. B) Cells transfected with expression plasmid encoding JY.8675309. C) Cells cotransfected with Ley2 and JY.8675309 expression plasmids. D) Cells cotransfected with expression plasmids encoding Ley2, JY.8675309, and short hairpin specific to Ley1 mRNA.

**Figure __.**
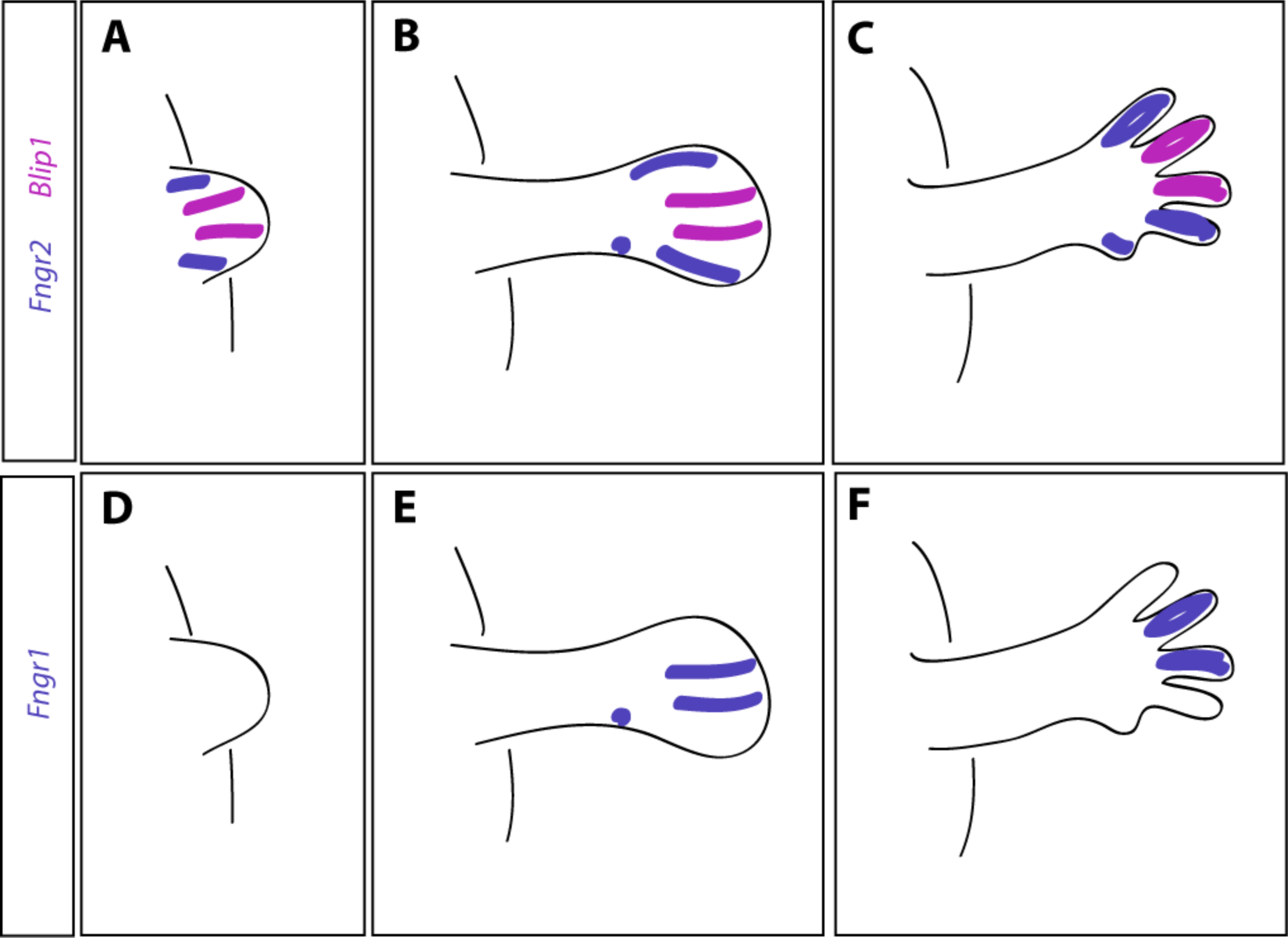
In situ hybridization detection of *Blip1* during limb development in the mouse. A) Expression of the new gene *Blip1* and known gene *Fngr2* at the limb bud stage. B) Expression of *Blip1* and *Fngr2* during limb bud outgrowth. C) Expression of *Blip1* and *Fngr2* in the fully formed limb. D) Expression of the known gene *Fngr1* at the limb bud stage. E) Expression of *Fngr1* during limb bud outgrowth. F) Expression of *Fngr1* in the fully formed limb.

**Figure __.**
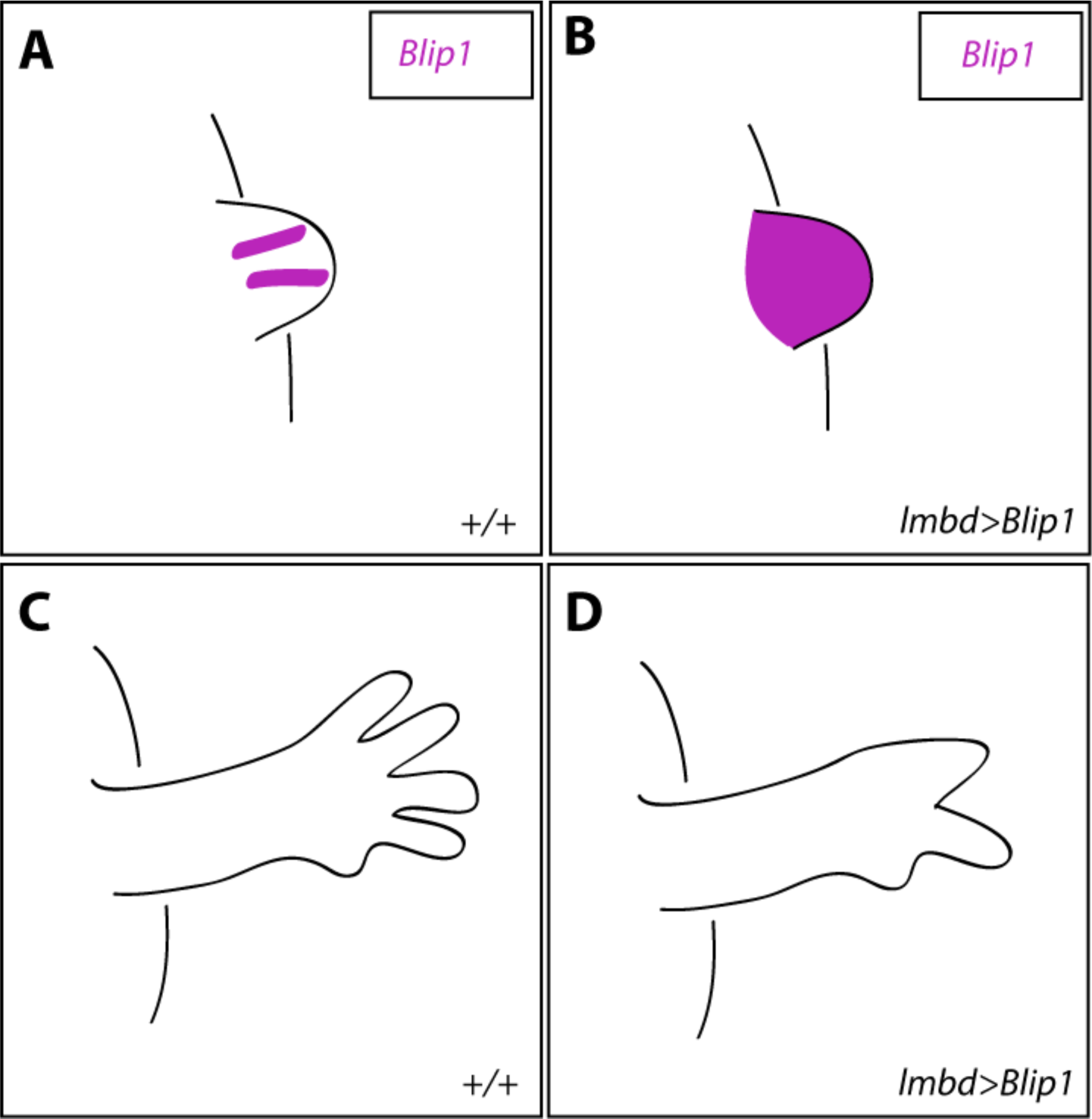
Phenotypic analysis of *Blip1* overexpression in the limb. A) In situ hybridization to show expression of the newly identified *Blip1* gene in the wild type limb. B) Expression of *Blip1* mRNA in limb buds following limb bud-specific overexpression of *Blip1.* C) Wild type mature limb. D) Mature limb resulting from genetic manipulation described in panel B.

**Figure __.**
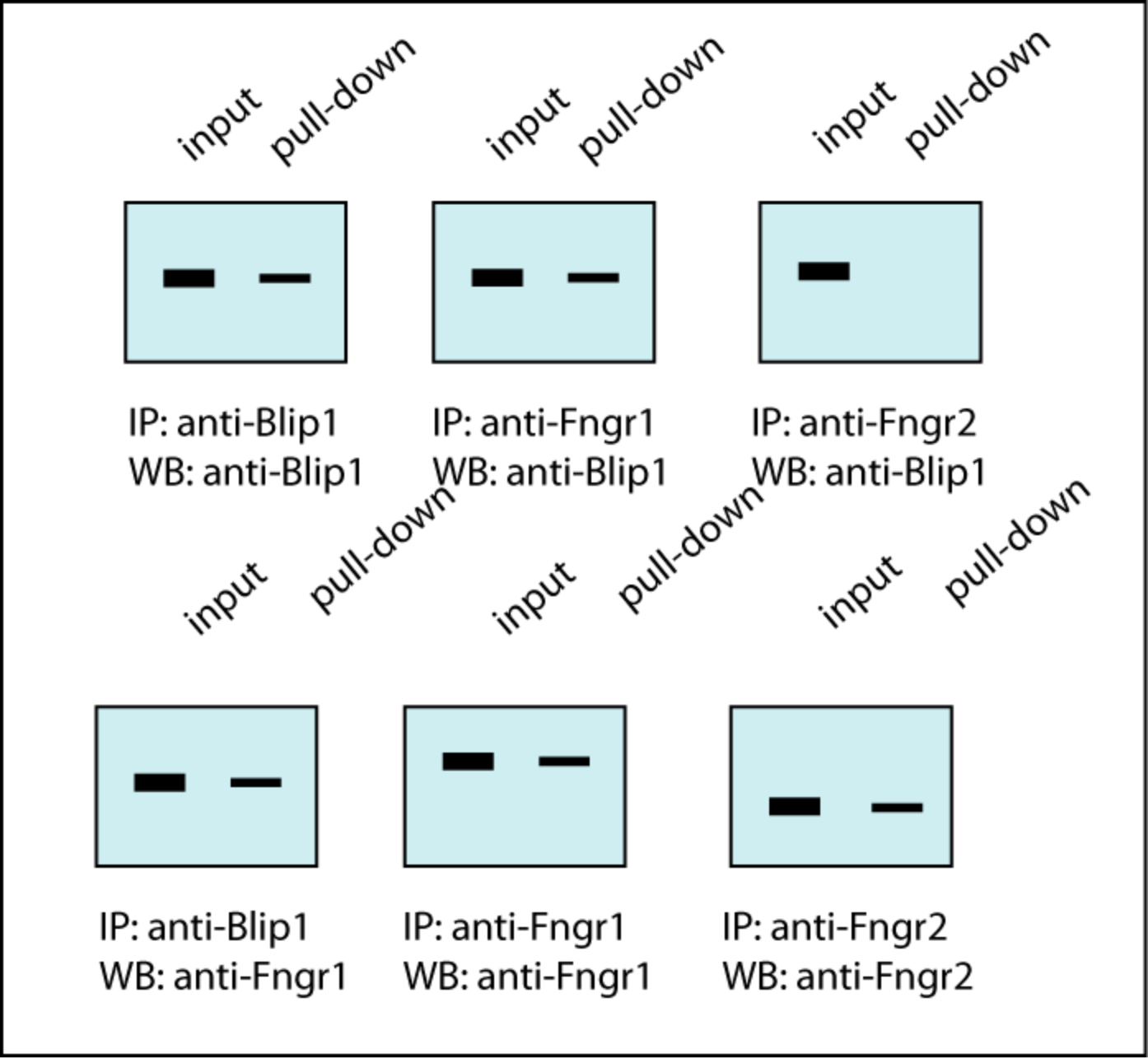
Coimmunoprecipitation of Blip1 and known limb patterning proteins. IP = antibody used for immunoprecipitation, WB = antibody used to probe western blot.

**Figure __.**
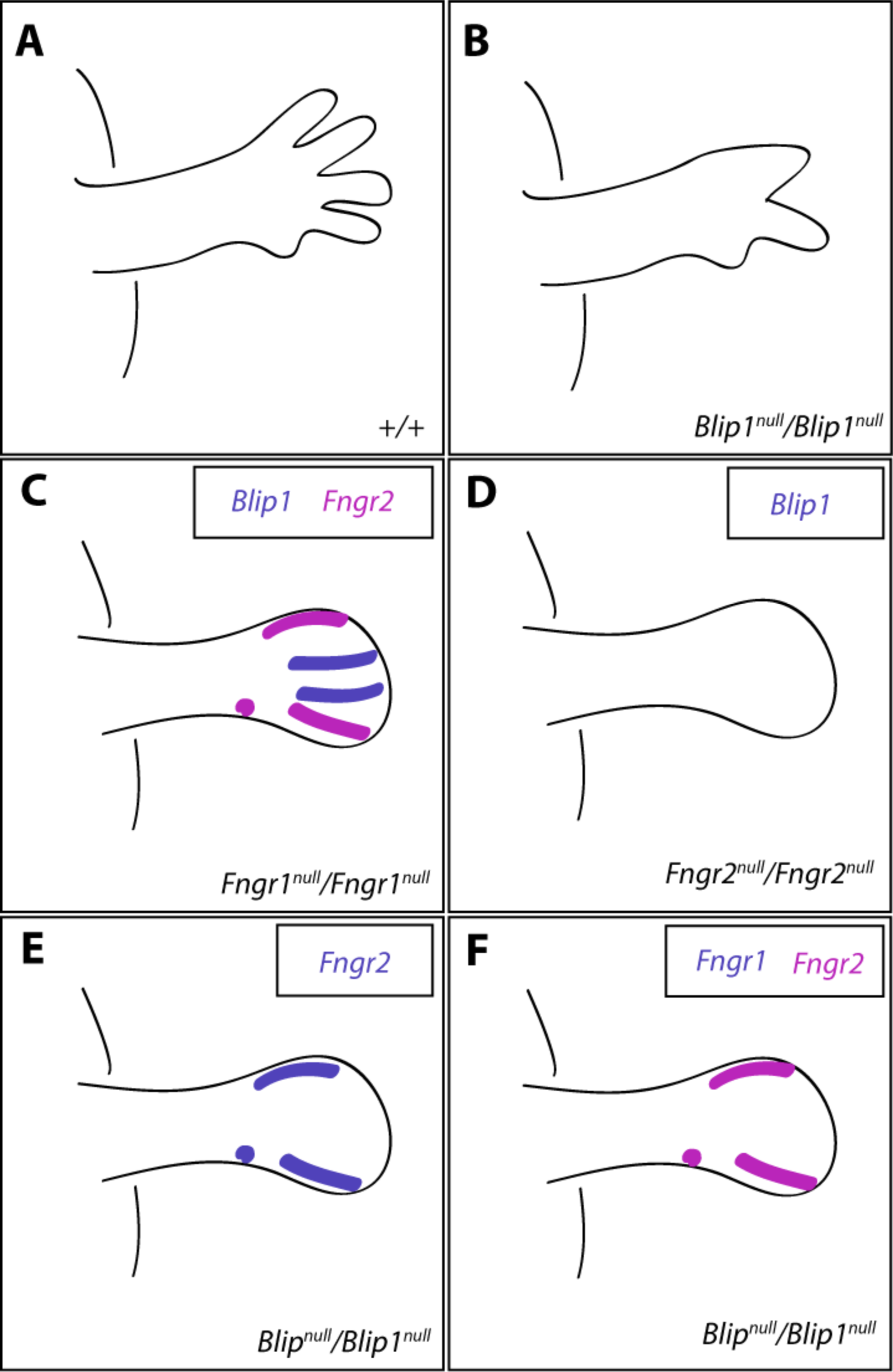
Phenotypic analysis of knockout mice lacking the new gene *Blip1* or known limb patterning genes *Fngr1* and *Fngr2*. A) Morphology of the mature wild type limb. B) Morphology of the limb in the *Blip1* knockout mouse, shown at the same stage as panel A. C) In situ hybridization to show expression patterns of *Blip1* and *Fngr2* mRNA localization in *Fngr1* knockout mouse during limb development. D) *Blip1* mRNA in *Fngr2* knockout mouse during limb development. E) *Fngr2* mRNA in *Blip1* knockout mouse during limb development. F) *Fngr1* and *Fngr2* mRNA in *Blip1* knockout mouse during limb development. Genes examined are color coded as indicated.

**Figure __.**
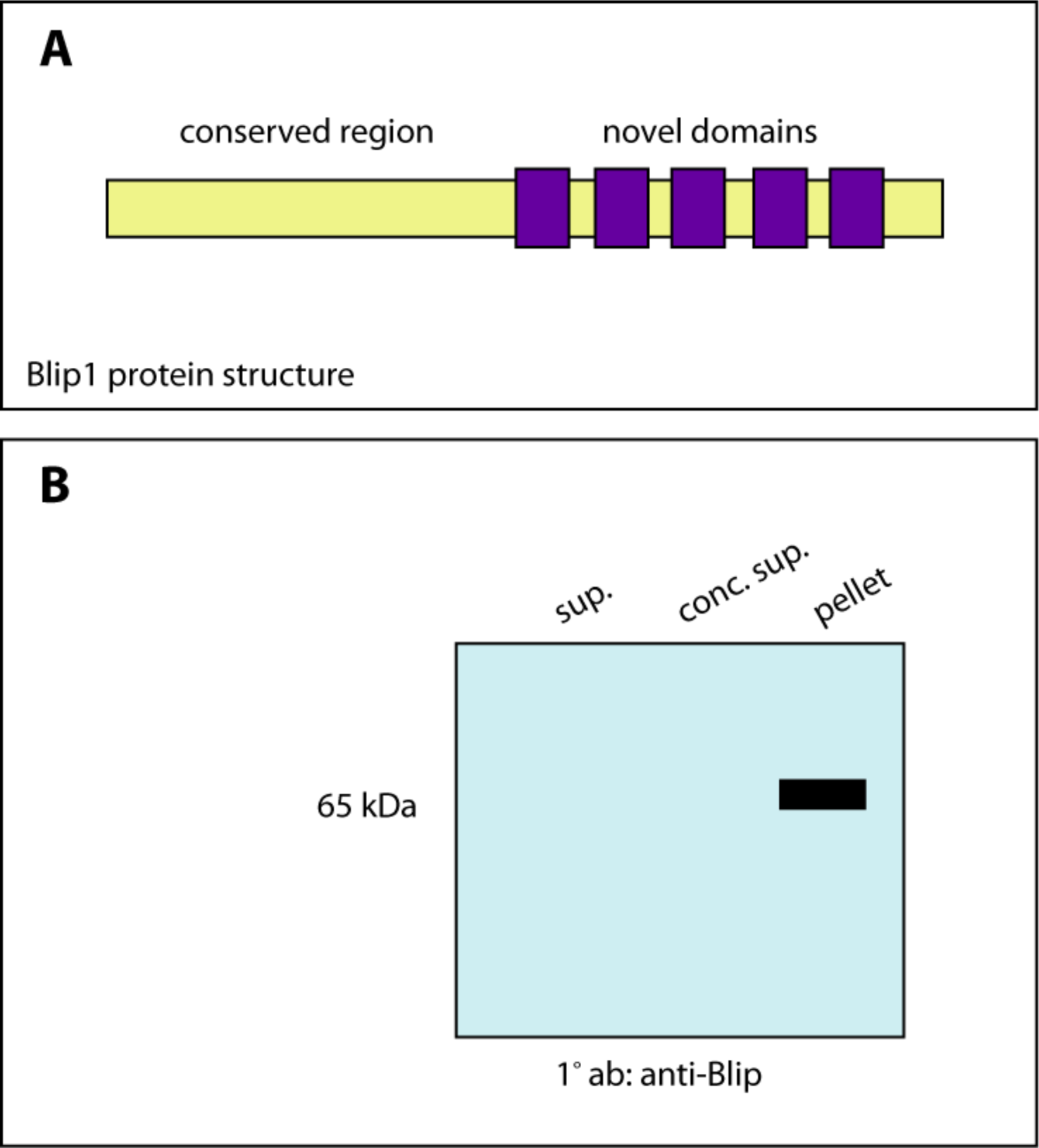
Biochemical characterization of Blip1 protein. A) Schematic representation of the Blip1 protein. B) Western blot to detect Blip1 protein in Hela cells transiently transfected with a Blip1 expression vector.

**Figure __.**
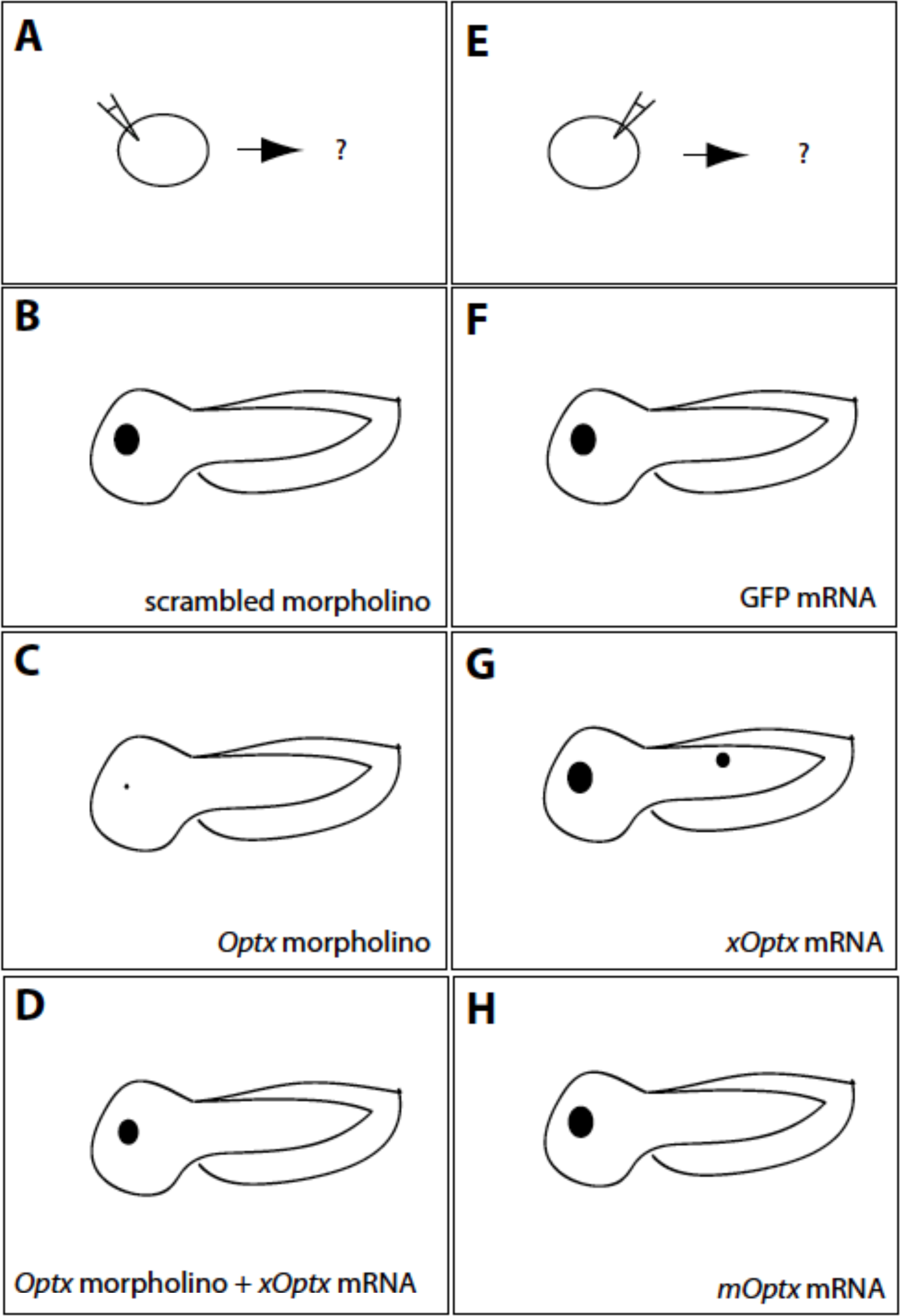
Manipula/on of levels of the newly identified *xOptx* gene during *Xenopus* development. A) Schematic overview of the experimental strategy leading to phenotypes shown in panels B-D. B) Larval phenotype following injection of a scrambled morpholino. C) Larval phenotype resulting from injection of *xOptx* morpholino. D) Larval phenotype resulting from coinjection of *xOptx* morpholino and mRNA encoding wild type *xOptx*. E) Schema*c overview of the experimental strategy leading to phenotypes shown in panels F-H. F) Larval phenotype resulting from injection of mRNA encoding *GFP.* G) Larval phenotype resulting from injection of mRNa encoding wild type *xOptx* mRNA. H) Larval phenotype resulting from injection of mouse *Optx* mRNA.

**Figure __.**
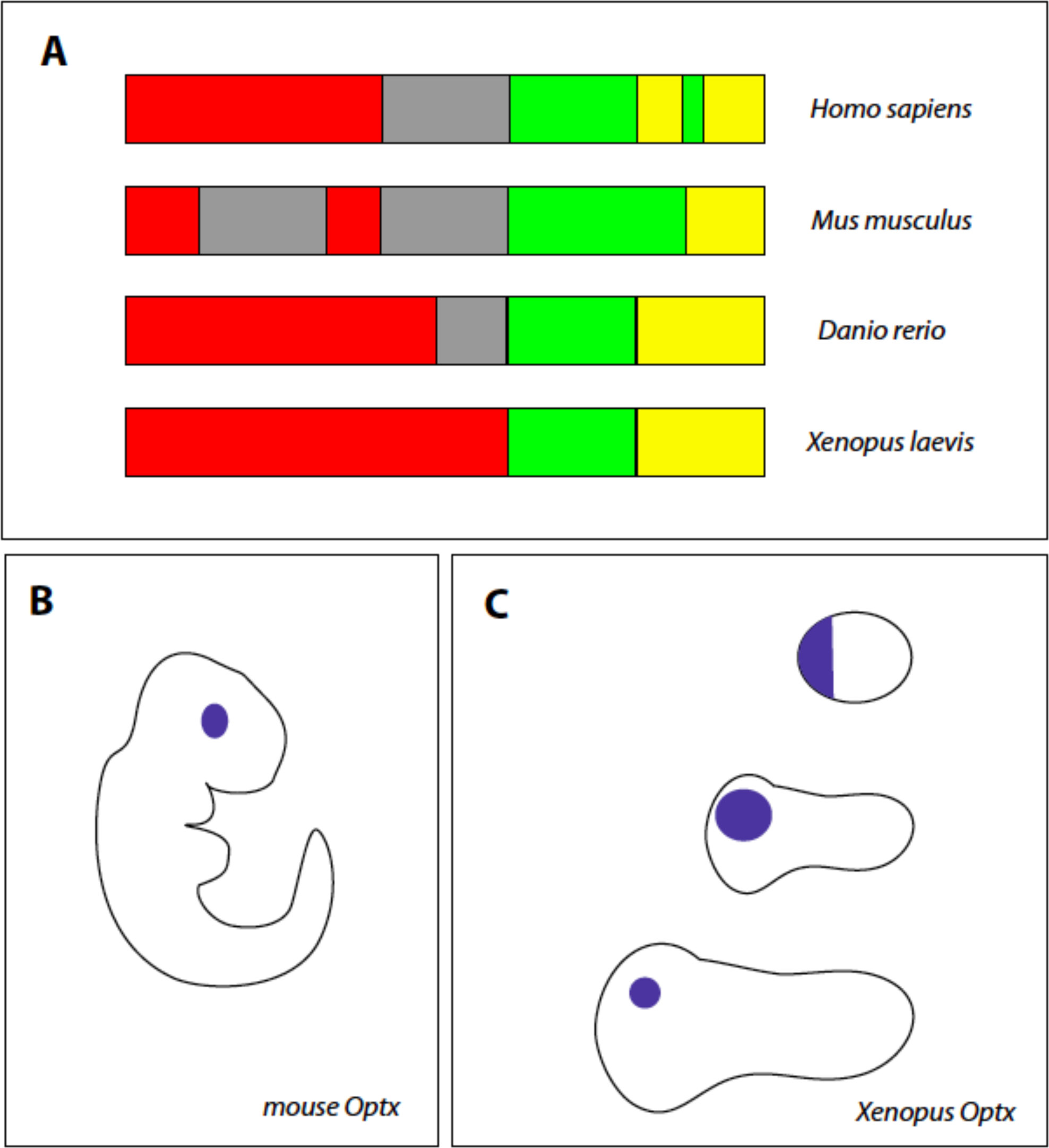
Characterization of the newly identified Optx protein in multiple species. A) Schematic representation of Optx proteins in indicated species. B) In situ hybridization of mouse embryos with the *mOptx* probe midgestation, prior to eye formation. C) Developmental time course of *xOptx* expression in *Xenopus* embryos prior to eye formation.

**Figure __.**
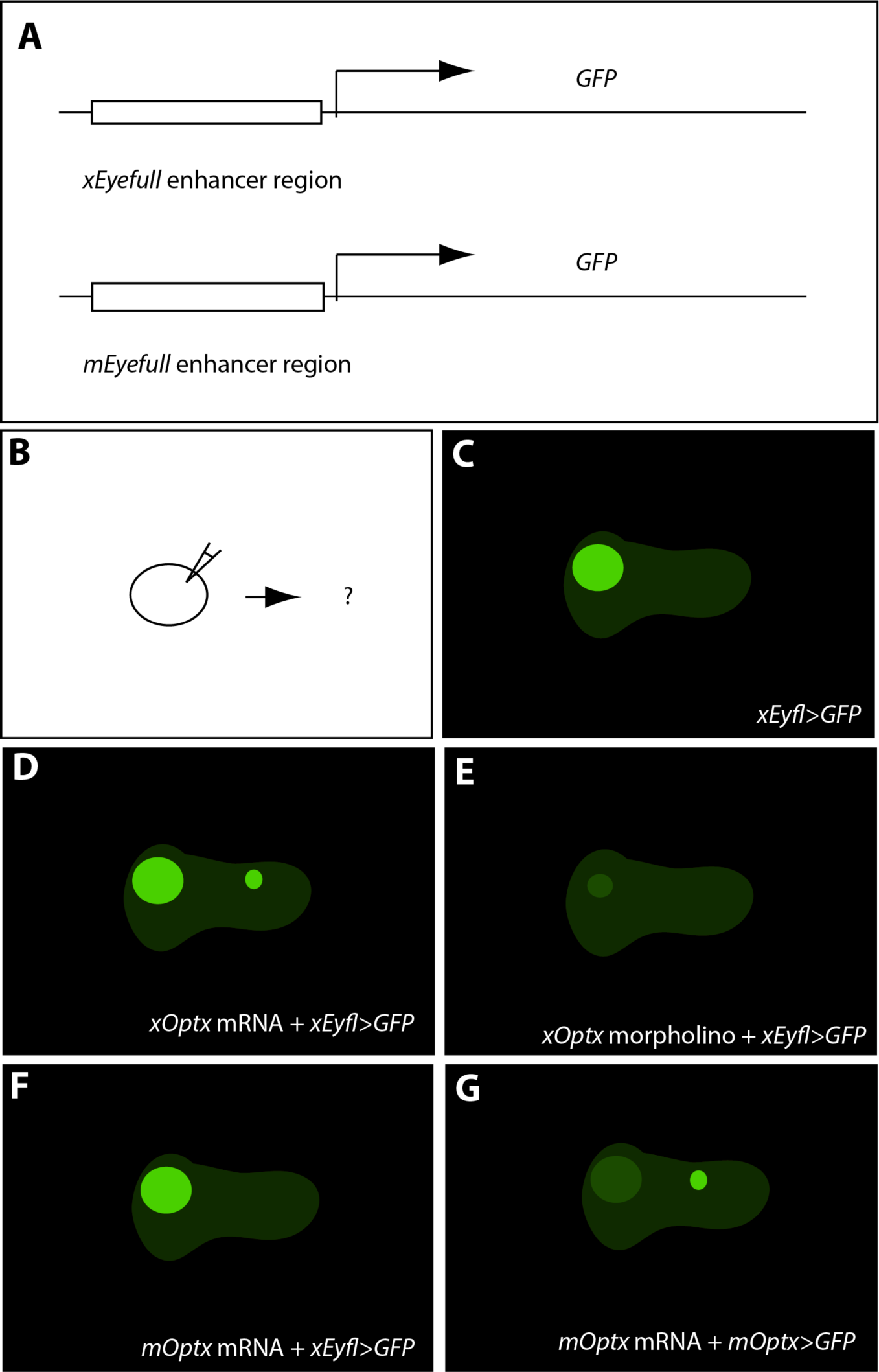
Regulation of the *Eyefull* (*Eyfl*) gene in the *Xenopus* embryo. A) Schematic representation of reporter constructs bearing *Xenopus* or *Musculus Eyfl* enhancer elements. B) Schematic representation of the experimental strategy leading to phenotypes shown in panels C-G. C) Larval GFP expression resulting from injection of mRNA encoding the *xEyfl* reporter. D) GFP following injection of *xOptx* mRNA and the *xEyfl* reporter. E) GFP following injection of *xOptx* morpholinos and the *xEyfl* reporter. F) GFP following injection of *mOptx* mRNA and the *xEyfl* reporter. G) GFP following injection of *mOptx* mRNA and the *mOptx* reporter.

## Homework 2 Instructions – You are the reviewer!

1. **Overview**

a. **Goal and how-to**
b. **Assessment**
2. **Reviewer Guidelines**

a. **Summary**
b. **Assessment Criteria**

1. **significancee**
2. **Observation**
3. **Interpretation**
4. **Model**
5. **Clarity**

### 1a. Goal and How-To

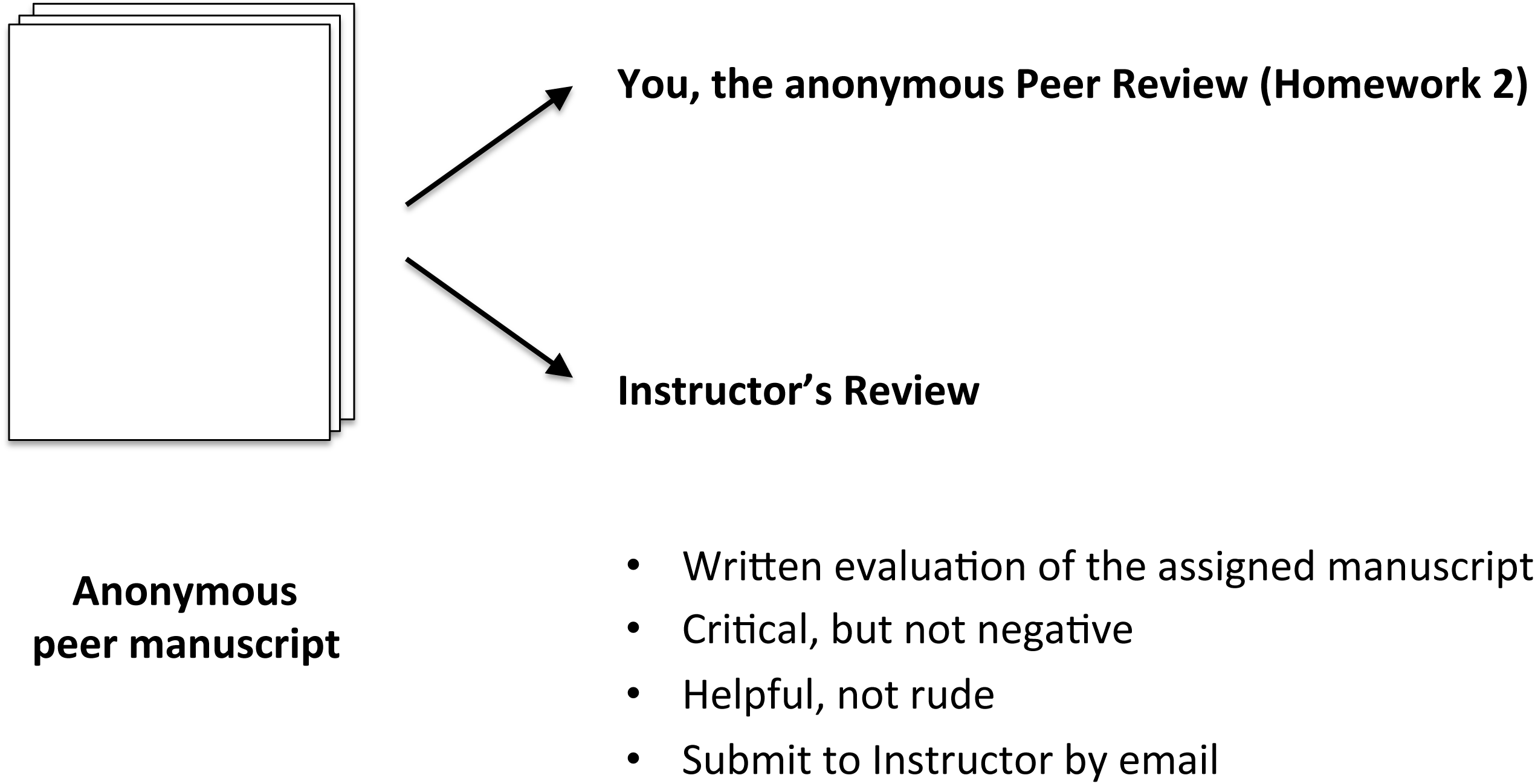

### 1b. Assessment

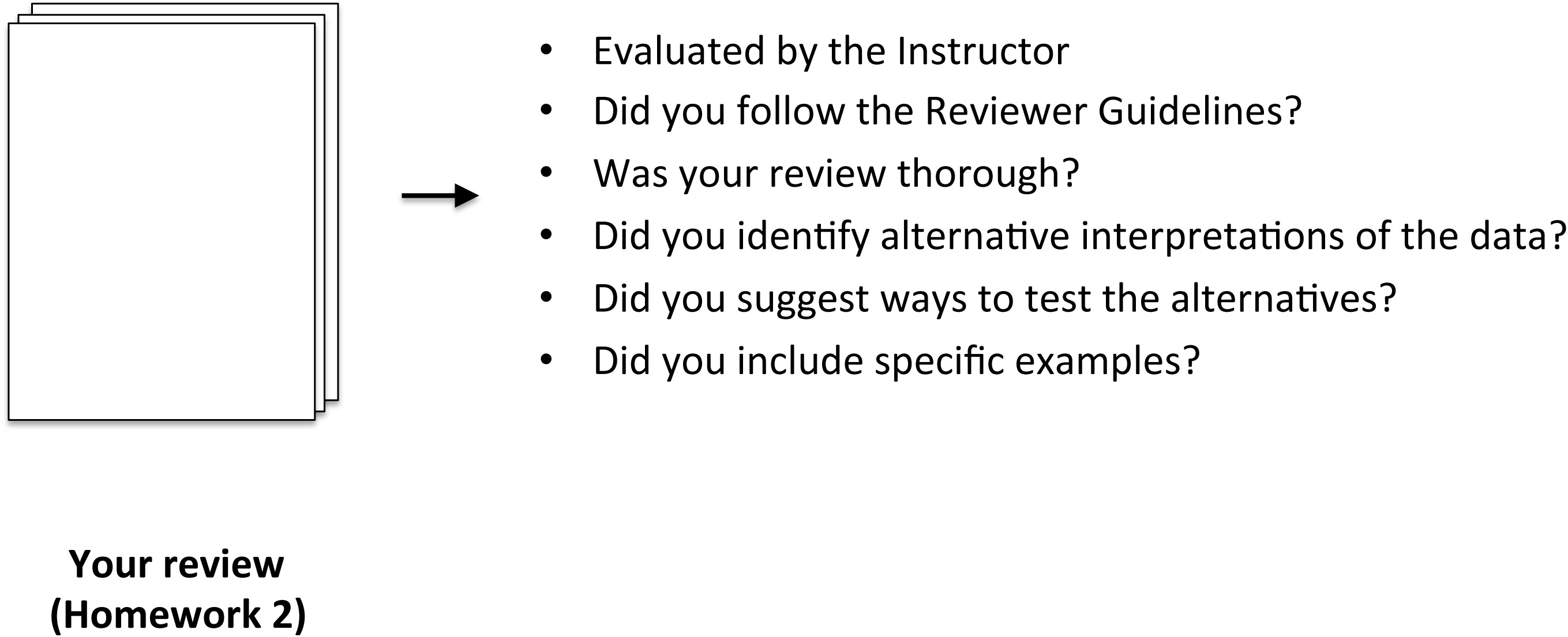

### 2a. Reviwer Guidelines - Summary

Limit: 300 words
- Describe the biological process explored in the manuscript
- Summarize the key findings of the manuscript
- Briefly describe the paper’s strengths and weaknesses, using the five assessment criteria (next slide)

### 2b. Reviewer Guidelines – Assessment Criteria

***Address each of the following with specific examples and suggestions for fixing the problem.***

1. **significancee** Do the authors explain the significancee of the study? Is their rationale for undertaking these studies compelling?
2. **Observation** Are the authors’ descriptions of the data accurate? Do they include a description of positive and/or negative controls?
3. **Interpretation** Are the authors’ interpretations supported by the observations? Are there alternative interpretations of the data that are not discussed? If so, what are they?
4. **Model** Is the authors’ molecular mechanism explained clearly in the text and in an original figure? Is the model supported by the observations? Did the authors state the predictions of the model and did they propose experiments to test the predictions of the model? Would the proposed experiments definitively test the model? Are there experiments that would be better?
5. **Clarity** Is the manuscript structured and formatted according to the *Author Guidelines*? Is the manuscript easy to read and free of jargon, typos, and grammatical errors?

